# The Effects of Inter-Individual Biological Differences and Taphonomic Alteration on Human Bone Protein Profiles: Implications for the Development of PMI/AAD Estimation Methods

**DOI:** 10.1101/2020.10.15.341156

**Authors:** Hayley L. Mickleburgh, Ed Schwalbe, Haruka Mizukami, Federica Sellitto, Sefora Starace, Daniel J. Wescott, David O. Carter, Noemi Procopio

## Abstract

Bone proteomics studies using animal proxies and skeletonized human remains have delivered encouraging results in the search for potential biomarkers for precise and accurate post-mortem interval (PMI) and the age-at-death (AAD) estimation in medico-legal investigations. At present, however, the effects of inter-individual biological differences and taphonomic alteration on recovered human bone protein profiles are not well understood. This study investigated the human bone proteome in four human body donors studied throughout decomposition outdoors. The effects of ageing phenomena (*in vivo* and post-mortem), and intrinsic and extrinsic variables on the variety and abundancy of the bone proteome were assessed. Results identified a new potential biomarker for PMI estimation, as well as three potential biomarkers for AAD estimation. The results also suggest that bone mineral density (BMD) may be an important variable affecting the survival and extraction of proteins.

**Highlights:**

- CO3, CO9, COBA2, CO3A1, MGP, PGS2 and TTHY are potential biomarkers for post-mortem interval estimation in skeletonized human remains
- FETUA, ALBU and OLFL3 are potential biomarkers for age-at-death estimation in human remains
- Taphonomic and biological variables play a significant role in survival and extraction rates of proteins in bone
- Bone mineral density may affect survival of proteins in bone, probably due to the effects of the mineral matrix on the movement of decomposer microbes
- Higher bone mineral density may affect the survival and the extraction rate of collagen and mineral-binding proteins

## 1. Introduction

Estimations of the time elapsed since death (post-mortem interval, PMI) and the age-at-death (AAD) are crucial in the forensic investigation of unidentified human remains. This information is important to distinguish between historical remains (>100 years old) and remains of medico-legal relevance (≤100 years old)^1,2^, and to narrow the search of missing persons for identification purposes^3,4^. High precision, accuracy and objectivity of PMI and AAD estimation methods are essential in order to be considered admissible in a court of law.

PMI estimation often relies on visual assessment of gross morphological changes of the body during decomposition^5–7^, even though the rate of these changes is known to be highly variable^8,9^. Accuracy of the PMI estimation decreases as decomposition progresses, and interobserver reliability differs depending on the method and the experience of the researcher^9,10^. Biochemical techniques have shown promising results in the search for a precise and accurate method to estimate late PMI in human bone, however, these methods are yet to be validated for use in forensic contexts^11–14^.

Standard AAD estimation methods are based on the examination of the morphological characteristics of the remains^15^, and require the evaluation of several different skeletal elements^16^. Different methods are applied to juveniles and adults^17,18^. Limitations of these methods include a high inter-observer variability^15^, inter- and intra-population variability with increasing AAD^19^, lack of consensus regarding the evaluation of the errors^20^, poor precision in adult aging in comparison with juvenile and adolescent aging^4^, and the requirement for remains to be as complete as possible^20^.

In recent years, bone proteomics methods have been demonstrated to be highly promising for the development of precise, accurate and objective PMI and AAD estimation methods, requiring only small samples of bone. Proteins are relatively stable in bone, and have been successfully extracted from archaeological^21–25^ and paleontological specimens^26–29^, making them a promising target for forensic applications^30^. Studies conducted using animal models (e.g., *Sus scrofa* and *Bos bovid)* focused on inter- and intra-individual comparisons and monitored changes in the bone proteomes associated with progressing decomposition stages. These studies revealed inter-skeletal proteomic variability^31^, and identified potential biomarkers for AAD^31^ and PMI estimations^32^. In addition, burial environment was found to affect the proteome recovered from archaeological specimens^33^. However, the development of bone proteomics methods for forensics remains impeded by the fact that it is unknown how representative animal models are for human specimens. Moreover, it is largely unknown how taphonomic processes and inter-individual variation (both *in vivo* and at the time of death), including underlying health conditions, affect the survival and extraction of bone protein profiles in humans.

A recent study conducted on human bones collected from a cemetery in Southeast Spain, provided promising new insights on the estimation of broad PMI ranges (5-20 years) in humans using protein biomarkers in proximal femoral bone^34^. The study identified 32 proteins which could be used in conjunction to discriminate between PMIs greater or smaller than 12 years^34^. The sampled individuals were subjected to similar taphonomic conditions, and PMIs were greater than 7 years in all but one case. While the study was conducted on a relatively large sample (n=40), inter-individual and inter-skeletal comparison of bone protein profiles at different stages of decomposition of the body were not possible as only one skeletal element was available per individual. For the further development, and ultimately validation, of forensic proteomics to estimate PMI, the study of changes in human bone protein profiles from the fresh stage of decomposition to the skeletonized stage is crucial.

In this study, we aimed to investigate the effects of taphonomy and biological variation on the recovery and variability of the human bone proteome, and evaluate potential avenues to develop a broadly applicable, standardized method of PMI and AAD estimation in human remains in advanced state of decomposition. The proteomes of anterior midshaft tibia and iliac crest samples from four body donors of known AAD (two buried and two placed in an open pit), taken shortly after death and upon complete skeletonization of the body, were analysed to investigate 1) whether the previously identified potential biomarkers for PMI and AAD are applicable to human bones with lower PMIs, 2) whether additional potential biomarkers for PMI/AAD estimation could be identified, 3) whether the human bone proteome is subject to inter-skeletal and inter-individual variability, and 4) the role depositional environment, season and taphonomy play in bone proteome recovery.

## 2. Results

### Proteomic data

The proteome of both the midshaft tibia and the iliac crest of four human body donors sampled at “fresh” (PMI = 2-10 days) and at “skeletonized” (i.e., when bodies did not have any adhering/desiccated soft tissue) stages of decomposition (PMI variable, between ~5200 and ~17800 ADD, depending on the season of placement, see Table 1), was analysed. Three replicate extractions were taken from each bone, totaling 48 proteomic analyses. After refining the Progenesis results based on the number of unique peptides and on the ion score (see Methods section), 133 quantifiable proteins were identified (Supplementary Data 1). The protein interaction network (Fig. 1) showed a significant enrichment of interactions (PPI enrichment *p*<1.0×10^−16^) and functional enrichments of specific GO terms for biological processes, cellular components and molecular functions (Supplementary Data 2).

**Figure 1.**
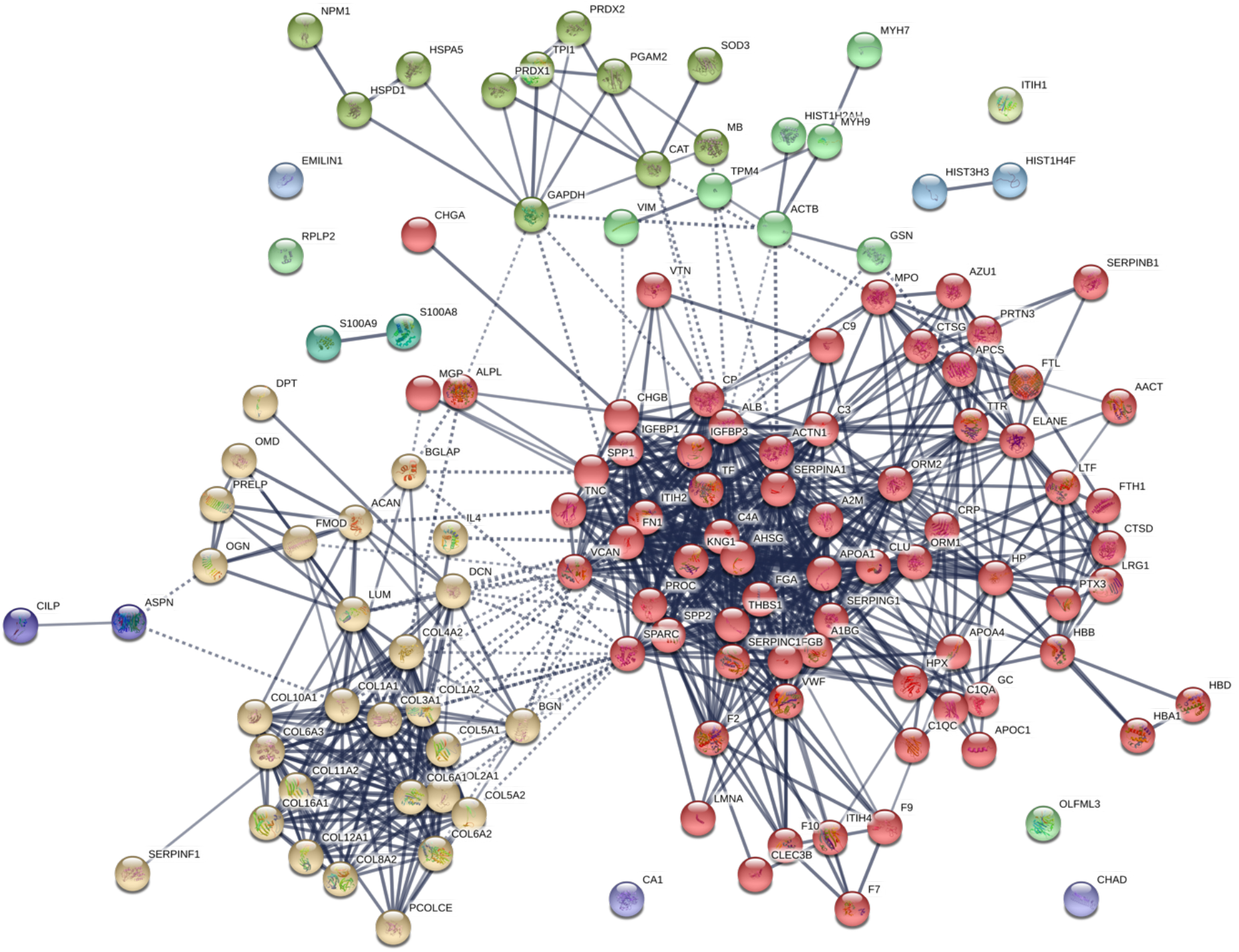
STRING protein network of the quantifiable proteins extracted from all samples. Immunoglobulin proteins (gene names IGHA1, IGHG2, IGHG3, IGKC, IGLC2) were not found with STRING and are not represented in the figure. The light orange cluster represents collagenous and collagen-binding proteins, the red one plasma and bone-related proteins, the yellow-green one at the top on the left side ubiquitous proteins and the light green one at the top on the right side some muscle proteins. Other smaller clusters represent other types of proteins interacting less with the major clusters identified.

**Table 1.**
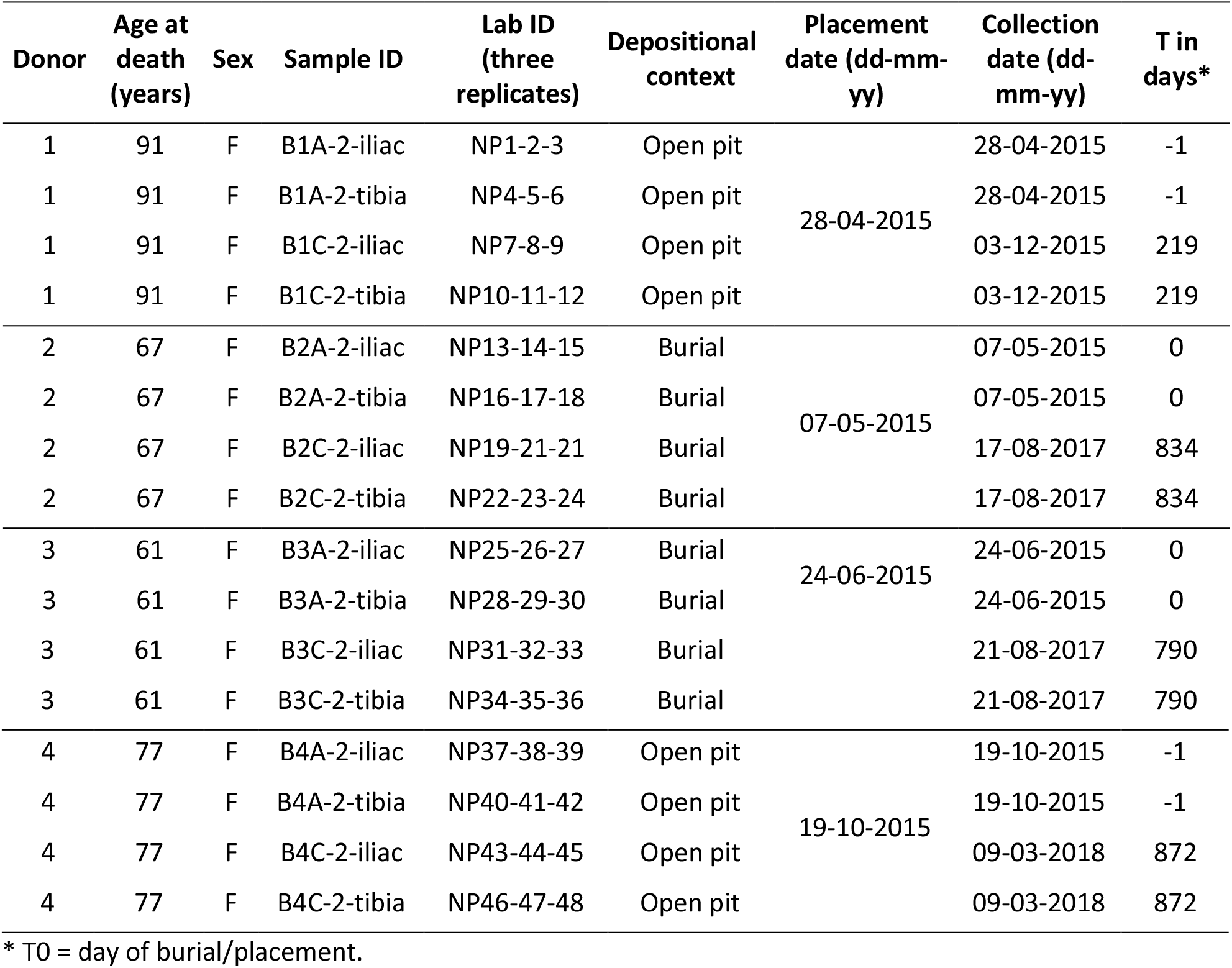
Biological and bone sample data.

### Human proteomic inter-skeletal and inter-individual variability

Fresh samples were found to have a significantly greater protein diversity than skeletonized samples (Fig. 2A), and fresh iliac samples were the richest samples analysed, both in terms of proteome diversity (average of 55 different proteins in iliac fresh samples versus 27 for tibia fresh, 15 for iliac skeletonized and 23 for tibia skeletonized, see Supplementary Data 3 for details) and protein relative abundances (Supplementary Data 4). In fact, among the 116 proteins with significantly different relative abundances between the various bone and sampling types (e.g., fresh vs. skeletonized), 105 (90.5%) were more abundant in the fresh iliac samples, eight (6.9%) in the skeletonized tibia samples, two (1.7%) in the fresh tibia samples and one (0.9%) in the skeletonized iliac samples (Supplementary Data 4). When comparing iliac fresh and skeletonized samples, 96 proteins had a significantly different abundance in the two groups, all of which were more abundant in fresh than in skeletonized samples (Fig. 3 and Supplementary Data 5). Comparison of the fresh and skeletonized tibia samples revealed 23 proteins with significantly different expression in the two groups, of which 19 were more abundant in the fresh samples and four in the skeletonized samples (Fig. 3 and Supplementary Data 5).

**Figure 2.**
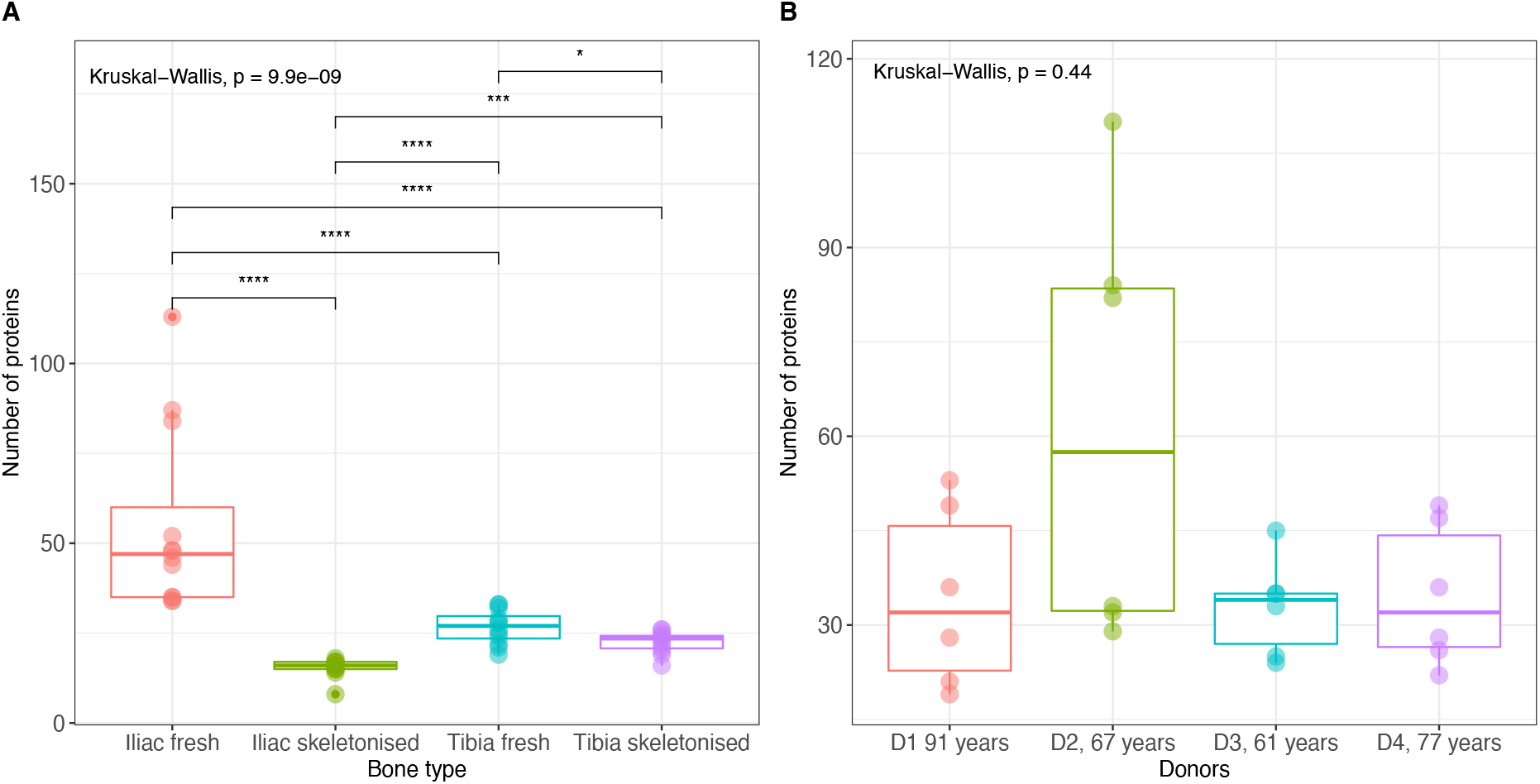
**A)** Number of proteins extracted from each sample. Samples were grouped according to bone type. All bone types were significantly different each other (post-hoc pairwise Wilcoxon-test with corrections for multiple testing, p value < 0.0002 = ****, p value < 0.002 = ***, p value < 0.01 = *). Outliers are represented as pointed-dots in the plot (two outliers identified here, one for iliac fresh (sample NP14, see Table 1 for details) and one for tibia iliac skeletonized group (sample NP45, see Table 1 for details)). **B)** Number of proteins extracted from fresh samples. Samples were grouped according to the donor. None of the donors resulted in being significantly different each other (post-hoc pairwise Wilcoxon-test with corrections for multiple testing, p value > 0.05).

**Figure 3.**
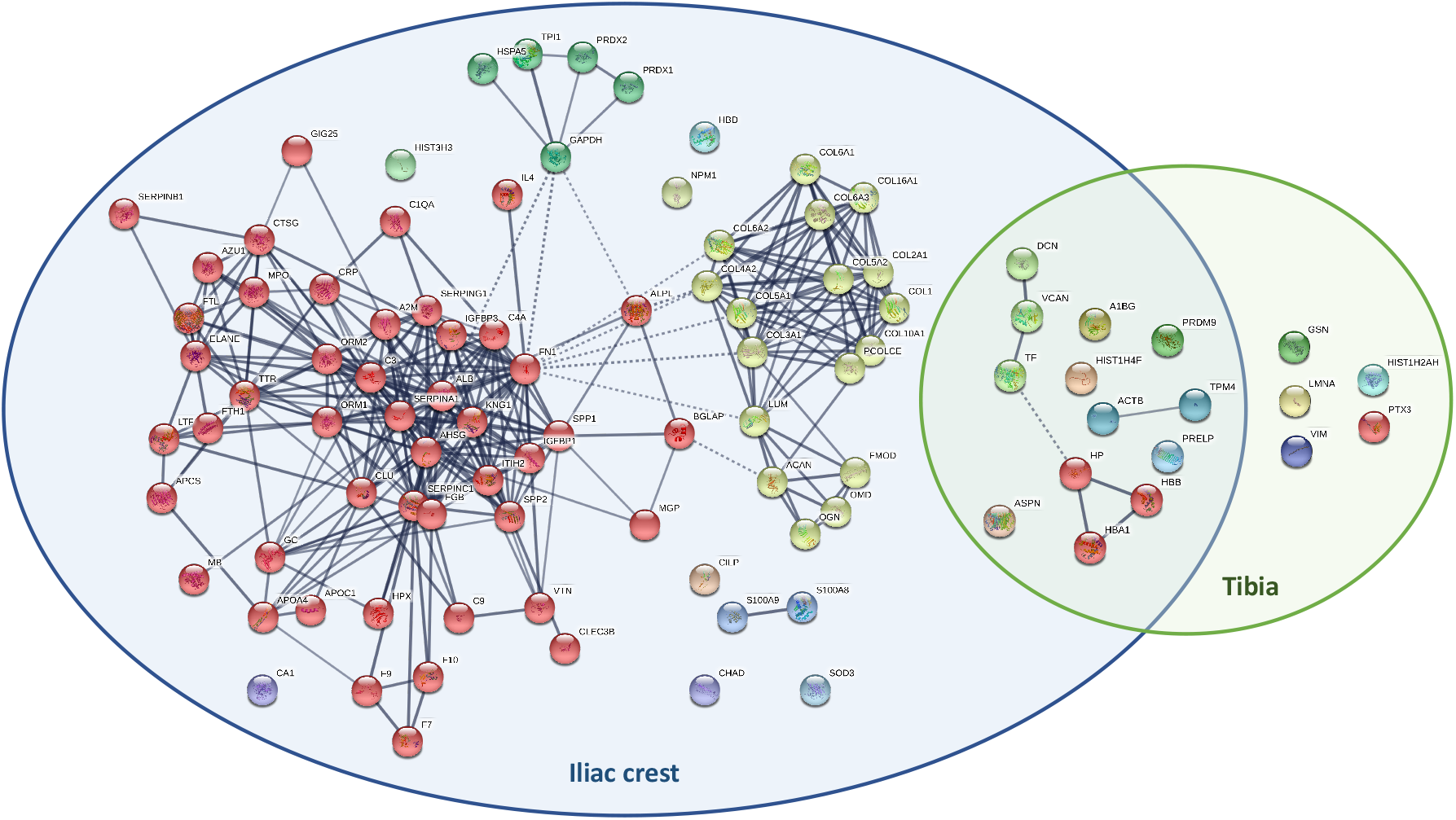
Venn diagram to represent STRING protein networks of proteins significantly more abundant in fresh iliac samples (left) and fresh tibia samples (right) than in their skeletonized counterparts. Proteins shared between the two categories are represented in the middle. Immunoglobulin proteins (gene names IGHA1, IGHG2, IGHG3, IGKC, IGLC2) were not found with STRING and are not represented in the figure. In the iliac crest category, red cluster represents plasma proteins, yellow cluster represent collagens and bone-related proteins, and green cluster ubiquitous proteins. No obvious clusters were identified for the shared proteins and for the ones belonging to the tibia category.

Comparison of inter-individual proteome variability of fresh bones only (to exclude any difference caused by taphonomic phenomena) showed that samples collected from D2 had a richer proteome variety (average number 62 for D2 versus 34, 33 and 35 for D1, D3 and D4 respectively), although this difference was not statistically significant (Fig. 2B). Comparing inter-individual relative protein abundances (Supplementary Data 6), we found 41 proteins that showed differences in their relative abundances among the donors. Of these, 36 (87.8%) were more abundant in D2, two each in D1 and D4 (4.9%) and one in D3 (2.4%).

### The influence of environment on bone proteome

Comparison of samples from the different depositional environments (open pits vs. shallow burials) showed no significant differences in the number of extracted proteins (p=0.3; Fig. 4A). Comparison of the relative protein abundances in these two groups revealed only four proteins with a different mean abundance for the two environments (three proteins more abundant in shallow burials, one protein more abundant in open pit placements, Supplementary Data 7). A test for association between the number of recovered proteins and the season of placement found no significant differences (p=0.4; Fig. 4B).

**Figure 4.**
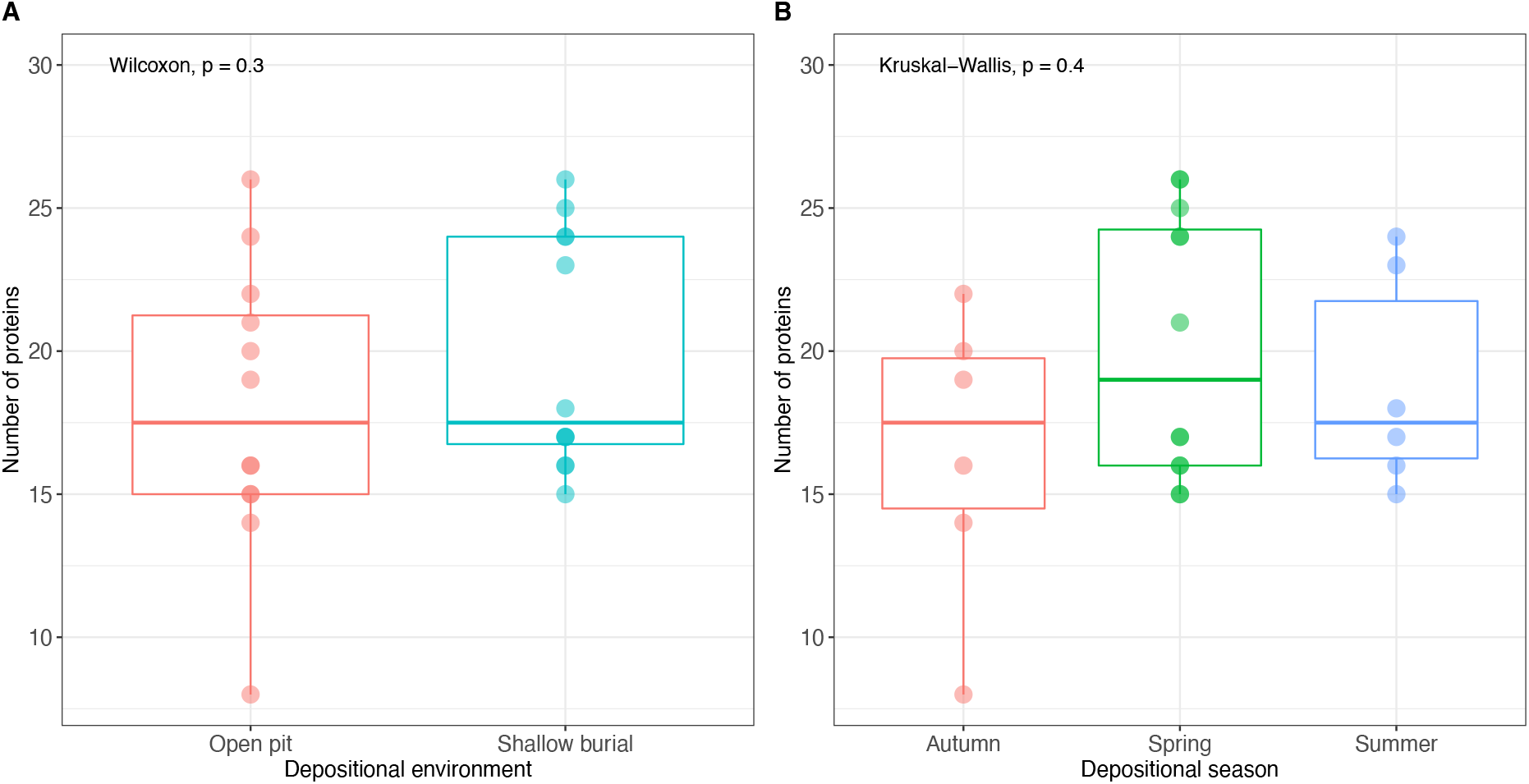
Number of proteins extracted from skeletonized samples, grouped by **A)** depositional environment or **B)** placement season. No significant differences were detected (Wilcoxon and Kruskal-Wallis p value > 0.05).

### Potential proteomic biomarkers for human PMI estimation

No association was found between the number of extracted proteins and the PMI of the samples. However, significant decreases in the abundance of collagen alpha-1(III) chain (CO3A1; p=0.0041), complement C9 (CO9; p=1.9e-06), collagen alpha-2(XI) chain (COBA2; 0.00055), matrix Gla protein (MGP; p=5.3e-05), decorin (PGS2; p=0.045) and transthyretin (TTHY; p=0.035) in iliac crest (Fig. 5A-F) and of complement C3 (CO3) in tibia (p=0.0.023; Fig. 5G) were observed when comparing the protein abundances of the skeletonized samples.

**Figure 5.**
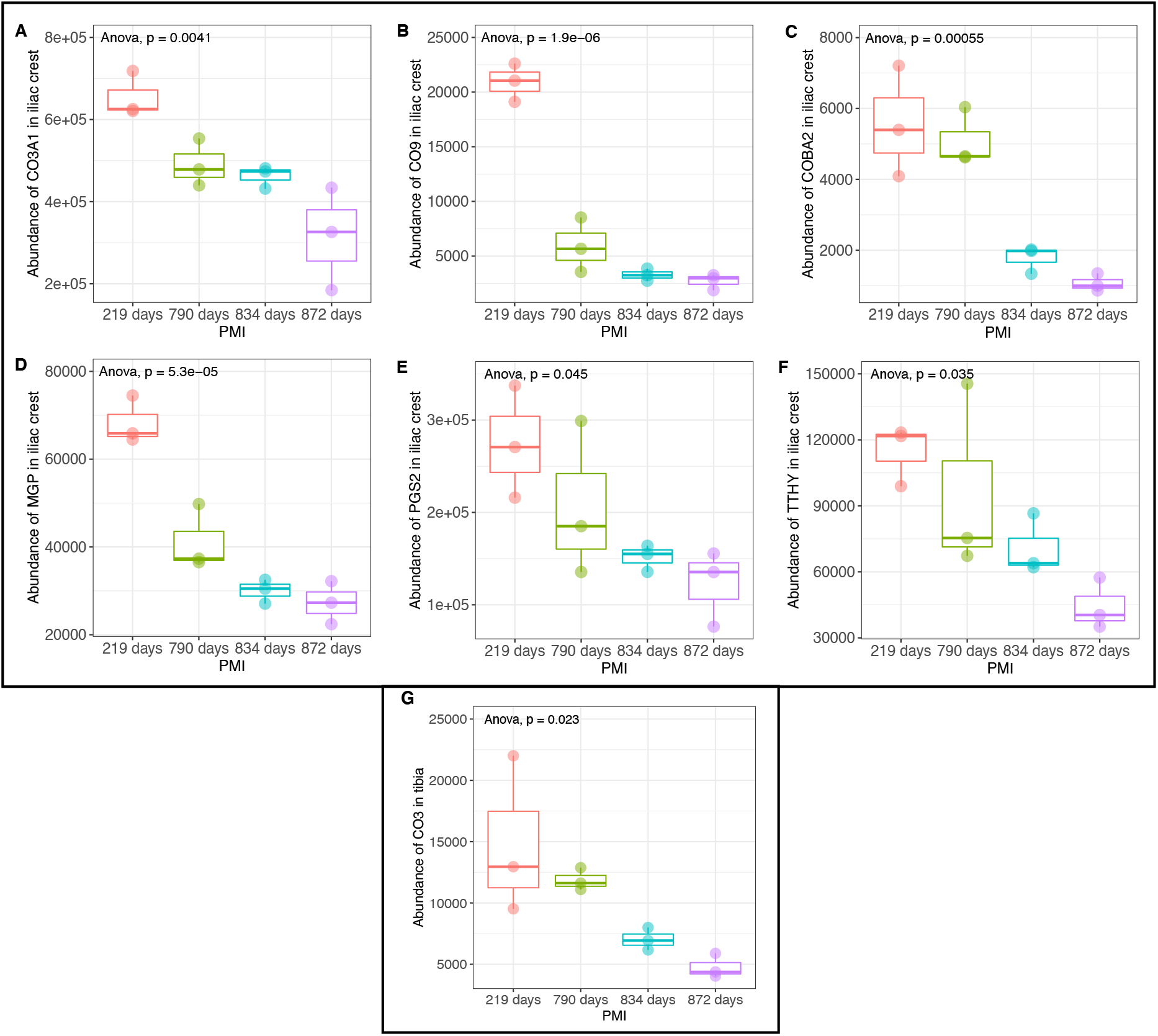
Abundance of **A)** CO3A1, **B)** CO9, **C)** COBA2, **D)** MGP, **E)** PGS2, **F)** TTHY protein in iliac crest skeletonized samples and of **G)** CO3 in tibia skeletonised samples with increasing PMIs. Groups are significantly different each other (ANOVA p value < 0.05).

### Potential proteomic biomarkers for human AAD estimation

The relative abundance of fetuin-A was found to be negatively associated with AAD in fresh tibia (p=0.033) and in skeletonized iliac samples (p=0.013). Skeletonized tibia samples showed lower levels for the oldest donor and higher levels for the other, but no significant trend was found (p=0.34). Iliac fresh samples showed similar levels in D2 and D3 and lower values for D1 and D4 (p=0.34; Fig. 6A-D).

**Figure 6.**
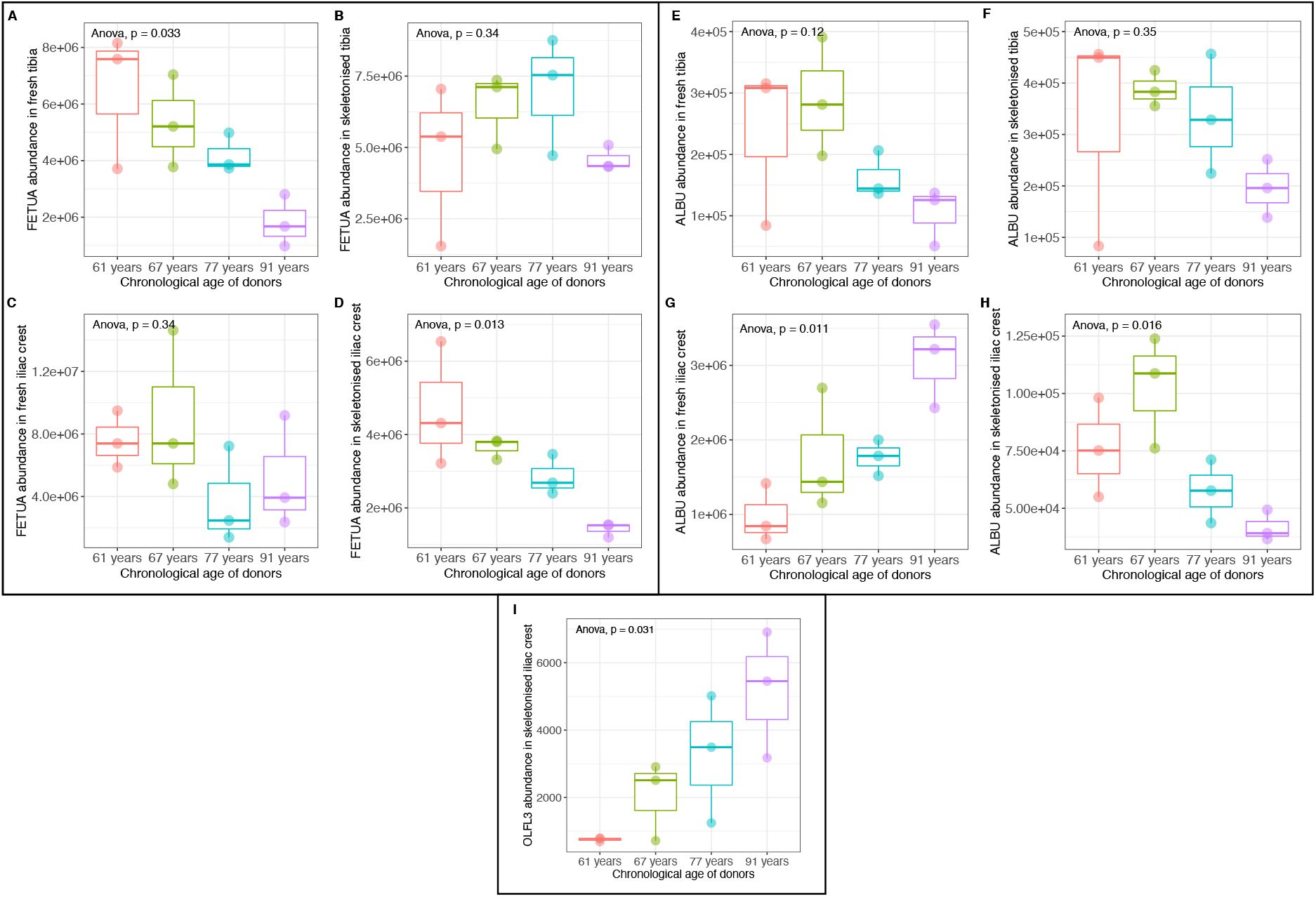
Relative abundance of fetuin-A in **A)** fresh tibia, **B)** skeletonized tibia, **C)** fresh iliac crest and **D)** skeletonized iliac crest samples, of albumin in **E)** fresh tibia, **F)** skeletonized tibia, **G)** fresh iliac crest and **H)** skeletonized iliac crest samples and of **I)** olfactomedin like-3 in skeletonized iliac crest samples, arranged by the chronological age of the donors. ANOVA p value was reported for each plot. Only A), D), G), H) and I) resulted in being statistically significant.

Significant differences in albumin abundance were found between different donors for both fresh (p=0.011) and skeletonized (p=0.016) iliac samples (Fig. 6E-H). In particular, fresh iliac samples showed a positive association with AAD, while fresh and skeletonized tibia samples both showed a negative relationship with AAD, although these results were not significant (Fig. 6; (p=0.12 and 0.35, respectively).

Additionally, a significant increase in the abundance of olfactomedin-like protein 3 (OLFL3) was observed in skeletonized iliac samples with increasing AAD (Fig. 6G; p=0.031).

## 3. Discussion

In this study, we identified specific proteins that significantly decreased in abundance with increasing PMI: complement C3 for tibia and collagen alpha-1(III) chain, complement C9, collagen alpha-2(XI) chain, matrix Gla protein, decorin and transthyretin for iliac crest. Four of the identified proteins are classified as bone structural/functional proteins (CO3A1, COBA2, MGP and PGS2) and three are plasma proteins (CO3, CO9 and TTHY). Previous work, conducted on animal proxies (pigs) left to decompose for a maximum of six months^35^, revealed a similar trend of consistently decreasing protein abundances over time, but for different proteins: haemoglobins, transferrins, triosephosphate isomerase, collagen alpha-2(V) chain and albumin. Both studies showed a reduction in the abundances of plasma and ubiquitous proteins, but reduction in bone structural/functional proteins was observed only in the current study. It is known that certain mineral-binding proteins (including structural ones) are susceptible to taphonomic processes of decay and diagenesis with prolonged PMIs^36^. The difference in duration and in the depositional environment and local climate between the pig study and the present study, and the resulting longer exposure to taphonomic processes, could therefore explain the different trends in mineral-binding protein abundances reduction that we observed. Analysis of human femoral bones from a cemetery context by Prieto-Bonete and colleagues^34^ also revealed a distinct reduction in the amount of structural and functional proteins in the highest PMI samples (13-20 years). Among the list of proteins identified in their study as biomarkers for prolonged PMIs, COBA2 was the only one that was also found here showing a similar inverse association with increasing PMIs in the present study. Overall, these findings suggest that COBA2 could be a good candidate for PMI estimation of human skeletonized remains, due to its durability over time and under different taphonomical conditions.

Our study found that the abundance of OLFL3, an osteoblast secreted extracellular matrix glycoprotein^37^, was positively associated with AAD in skeletonized iliac samples, adding a previously unreported protein to the list of potential biomarkers for AAD. Previous studies on animal and archaeological human bones identified a negative correlation between serum fetuin-A and AAD^21,38^, and proposed fetuin-A as potential biomarker for AAD^39^. The present study identified a similar negative relationship in fresh tibia and skeletonized iliac crest samples, but not in fresh iliac and skeletonized tibia. In addition to fetuin-A, several studies showed that serum albumin concentration is negatively correlated with AAD^40,41^. The present study found a non-significant negative correlation both in fresh and skeletonized tibia samples, but an opposite and significant trend in fresh iliac samples. Considering the relatively small sample size, it is difficult to interpret these findings. Based on observations of bone properties (hardness, weight, coloration) during sampling of the bone we postulate that inter-individual differences in bone mineral density (BMD; discussed below) as well as the inter-skeletal differences in BMD (i.e., iliac bone is less densely mineralised than tibia), may have affected these results. BMD is known to vary between different parts of the skeleton^42^. With regards to inter-individual BMD differences it is important to note that our observations do not constitute quantified measurements of mass-to-volume ratio, but are observations of the physical properties of bones during drilling. Our study used three biological replicates, however using a larger number in future studies might clarify whether fetuin-A and albumin are consistently negatively correlated with AAD in humans in different bone types and therefore can be used as a biomarker for AAD.

The potential effects of BMD on the recovered protein profile are suggested by the results of inter-individual comparisons. Fresh tibia samples from all four donors showed greater inter-individual reproducibility than fresh iliac samples. Fresh iliac samples from D2 showed significantly increased protein variety and abundances. Observations during drilling of the bone samples suggest that BMD in D2 may be relatively high. BMD could theoretically affect the variety and abundance of specific non-collagenous proteins that can either bind the calcium ions or the collagen in the mineral matrix^26^, thereby affecting the overall protein profile. In addition to non-collagenous proteins, many ubiquitous and plasma proteins were identified predominantly in fresh iliac samples, particularly from D2. Due to the greater blood irroration of the iliac crest in comparison with the midshaft tibia, a greater variety of these types of proteins could be expected in fresh iliac samples (discussed below). In addition to the observations of bone hardness in D2 made during drilling, the medical information available for D2 indicates certain conditions and treatments received in the years prior to death that are associated with changes in BMD, including chemotherapy treatment for cancer, prolonged consumption of calcium lactate^43^ and possible use of probiotics as adjuvant during cancer treatment^44,45^.

The likely greater BMD of D2 may have allowed for a stronger in vivo embedding of these proteins within the mineral matrix, resulting in the greater proteomic variety observed here. This effect of BMD on protein linkage in bones would be analogous to positive relationships observed between organic matter content and soil density, which is often a function of clay content. In fact, clay particles tend to carry a negative charge to bind with nutrient cations such as calcium and potassium, and these bonds can protect proteins from decomposition and even from extreme environmental conditions such as autoclaving^46^.

The results of inter-skeletal comparisons of the skeletonized samples suggest that both BMD and taphonomic processes affected the preservation and extraction of the bone proteome. The highly irrorated and less densely mineralized fresh iliac samples yielded greater variety and abundance of proteins, including those expressed specifically in plasma. The proteomes recovered from skeletonized iliac samples demonstrated significant protein decay occurred in this bone. The denser and less irrorated fresh tibia samples yielded lower protein variety and abundances by comparison to the fresh iliac samples. Comparison of the fresh tibia samples with the skeletonized tibia samples showed protein decay also occurred in this bone, but not to the degree observed in the iliac crest. These results suggest that differences in BMD and blood irroration between the iliac crest and the midshaft anterior tibia affected both the successful extraction of proteins from fresh samples, as well as the preservation in and extraction of proteins from skeletonized samples. Taphonomic processes of decomposition are known to affect BMD in humans, and can differentially affect skeletal elements^47,48^. Higher porosity of the iliac exposed this bone to significant deterioration as a result of taphonomic processes over time, resulting in the reduced inter-individual differences observed in the skeletonized iliac samples. Protein extraction from the dense and poorly irrorated fresh anterior midshaft tibia was less successful – at least with the mild extraction protocol used here – but the structure of this bone led to less taphonomic deterioration over time.

The current results likely indicate the effects of the decomposer community and physicochemical environment on the decomposition of human remains. A less dense matrix would facilitate leaching while promoting the movement of decomposer microbes throughout the bone. Microbial induced bioerosion, which is characterized by the chemical dissolution of mineral components of bone followed by the microbial enzymatic attack of organic components of bone, is thought to be one of the main causes of bone diagenesis^49^. The movement of decomposer microbes might be restricted to the external surfaces of more densely mineralised bone. The effect of the decomposer microbial community may be further influenced by the location of the bones. The iliac crest might be subject to greater decomposition because it is located closer to the trunk, which contains a significant amount of moisture and a large gastrointestinal microbial community that is known to translocate during decomposition^50,51^. The iliac crest, therefore, is likely located in a microhabitat that is favourable for decomposition. In contrast, the tibia is located further from the trunk in limbs that often desiccate during decomposition. The body position during decomposition of the four donors was flexed, and allowed the anterior tibiae to remain elevated above decomposition fluids excreted from the trunk. Desiccation of the soft tissues around the anterior tibiae was observed early on during decomposition of both open pit placements. Desiccated, densely mineralised bone is unlikely to be favourable for decomposition.

Comparison of samples from open pit placements with samples from burials, as well as comparison of season of placement in this study found no significant differences. While archaeological remains have revealed differences in protein recovery related to depositional environment^23,26,33^, it is possible that due to the relatively short duration of this experiment such environmental effects were not measurable in this study. It is also possible that the two depositional environments did not produce distinct enough conditions (Supplementary Table 1) to cause noticeable differences in the preservation of the biomolecules.

The preliminary indications from this study support previous findings that specific proteins decay at different rates, strengthening the potential for developing bone proteomics PMI estimation methods. COBA2 appears to be a good candidate for PMI estimation of skeletonized remains, together with CO3A1, PGS2 and MGP. The blood proteins CO3, CO9 and TTHY may be good candidates for shorter PMI estimation (i.e., before the complete degradation of blood proteins). Our study only partially supported previous studies identifying fetuin-A and albumin as potential biomarkers for AAD estimation, and additionally found OLFL3 being positively correlated with AAD. At the same time, our findings suggest that taphonomic (e.g., microbial bioerosion) and biological (e.g., variation in BMD) variables play a significant role in both the survival and extraction rate of proteins, due to their effects on the protective mineral matrix.

While the sample size is relatively small, the findings point toward potentially significant effects of inter-individual variation associated with health conditions, medical treatment, and possibly food and supplement intake. The results of both inter-individual and inter-skeletal comparisons in our study suggest that BMD may be an important variable affecting the survival and extraction of proteins in the bone mineral matrix. Higher BMD may promote attachment of a greater abundance and variety of mineral binding proteins, and in highly irrorated bones may additionally help to preserve more plasma proteins within the mineral matrix. Inter-skeletal differences in BMD appear to lead to distinct differences in the variety and abundance of preserved (and extracted) proteins. The attachment of proteins within a more densely mineralized bone matrix may protect them during microbial bioerosion. Based on these indications, we recommend including standard measurement of BMD and targeting a combination of different biomarkers (i.e., abundances of selected plasma proteins and bone-specific proteins) in future work. Overall, these results emphasize the limitations of developing methods and models based on animal proxies, since farmed animals rarely show the degree of inter-individual dietary and activity related variation that humans do, and BMD and degree of irroration of bones differ between species^52^. Moreover, these results emphasize the importance of conducting replication studies in larger human samples representing a broader range of PMIs and AAD, as well as sampling different bones, to better understand how different types of proteins and different parts of the human skeleton are affected by inter-individual variation and taphonomic processes. Finally, preliminary evaluation of the inter-skeletal differences we observed suggests that for future development of proteomics PMI estimation methods, the iliac crest bone may be a more suitable sampling target for relatively fresh remains of forensic interest and for archaeological studies specifically targeting the serum-proteins, due to the presence of greater protein variety of bone-marrow proteins. Specific burial conditions, such as dry burial environments, anaerobic environments, and certain post-mortem treatments of the body (such as embalming procedures) can limit the amount of bone diagenesis^53,54^ thereby promoting the survival of bone proteins across archaeological timeframes. In such circumstances the iliac crest may provide better results than the tibia to detect pathologies and infections associated with the bone marrow. The midshaft tibia may be a more suitable sampling target for skeletonized remains or those in a state of advanced decomposition of forensic interest, due to the better survival of collagen and mineral-related proteins that could be ultimately used for developing new biomolecular methods for PMI/AAD estimation.

## 4. Methods

### Body donations

The body donations of four females aged between 61 and 91 years old were placed unclothed to decompose at the Forensic Anthropology Research Facility (FARF), the outdoor human decomposition facility associated with the Forensic Anthropology Center at Texas State University (FACTS), between April 2015 and March 2018. While the targeted bone proteins in this study are not thought to differ between males and females, only post-menopausal female individuals were included, in order to exclude biological sex and major hormonal differences as a potential variable from the study. Two body donations (D2 and D3) were buried with soil in shallow hand dug pits. Two body donations (D1 and D4) were placed in pits of similar dimensions, which remained open throughout the experiment. Open pits were covered with a metal cages to protect the remains from large scavengers. The sample size in this study reflects general trends in human decomposition research, in which larger samples – like those used in clinical studies – can be difficult to obtain for practical, logistical and ethical reasons. While animal analogues such as pigs can be used to alleviate some limitations associated with small sample sizes, the study of human cadavers is important due to biological differences between humans and pigs, including anatomical differences in the digestive vasculature, and molecular differences in adipose tissue^55^.

Data on body decomposition and weather were collected throughout the experiment and can be found in Supplementary Table 1. Additional information on FARF’s environment can be found in the Supplementary Information. Gross decomposition was quantified using the total body score (TBS) method following Megyesi et al.^56^. Accumulated degree-days (ADD) were calculated using temperature data recorded on the facility.

### Bone sample collection

Bone samples (ca. 1 × 1 cm) of the anterior midshaft tibia and iliac crest (left) were collected prior to placement of the fresh body outside and upon retrieval of the completely skeletonized remains (right). The total 16 bone samples were stored in sterile plastic bags, and immediately transferred to a lockable freezer at −80ºC. Samples were shipped overnight on dry ice to the to the Forensic Science Unit at Northumbria University, U.K. Upon arrival, the samples were immediately transferred to a lockable freezer at −18ºC, adhering to the U.K. Human Tissue Act under the license number 12495. The experiment was reviewed and approved by the ethics committee at Northumbria University, with the reference code 11623. All biological and bone sample data are provided in Table 1. Observations on bone condition (density and colour) during sampling can be found in the Supplementary Information.

### Sub-sampling and sample preparation

The 16 samples were defrosted prior to their analysis, then cleaned in deionized water for three hours at room temperature, exchanging the water three times, once every hour. They were then dried in a fume cupboard at room temperature until completely dry. Bone samples were then secured in a table clamp for the sampling. Contamination between samples was prevented by using a double layer of aluminium foil within the clamp (in contact with the bone) and by using new foil double layers for each piece of bone sampled. The clamp was also cleaned in between each sampling step using 50% sodium hypochlorite (Sigma-Aldrich, U.K.), to further prevent contamination issues. Once the bone was secured in the clamp, Dentist’s Protaper Universal Hand Files (Henry Schein Minerva Dental, U.K.) were used to hand-drill ~25mg of fine bone powder for three times (i.e. three samplings were performed on the same bone fragment), in order to obtain three replicates for each of the bones analysed. By sampling in different locations close together on the same bone, we obtained multiple biological samples. Since it is known that bone proteins can vary throughout the human skeleton and within individual bones, these biological replicates, in contrast to technical replicates, allow us to assess the degree of intra-bone variability and to establish whether inter-individual differences are greater than the intra-bone variability, as indicated in a previous study using pigs as proxies^31^. Protaper files were changed between each sample, to prevent contamination. When the bone samples were too porous to obtain a fine bone powder (e.g., iliac crest samples), small bone fragments were cut using the Protaper files, and ~25mg of bone fragments were collected for each of the three subsamples in order to have three replicates.

### Protein extraction

Overall, 48 samples were obtained from the 16 bone pieces, and were subjected to bone protein extraction following the protocol of Procopio and Buckley^57^.

Briefly, each sample was decalcified with 1 mL of 10 v/v% formic acid (Fisher Scientific, U.K.) for 6 hours at 4 °C. After removing all the acid soluble fraction, the acid insoluble fraction was incubated for 18 hours at 4°C with 500 μL of 6 M guanidine hydrochloride/100mM TRIS buffer (pH 7.4, Sigma-Aldrich, U.K.). The buffer was exchanged into 100 μL of 50 mM ammonium acetate (Scientific Laboratory Supplies, U.K.) with 10K molecular-weight cut off filters (Vivaspin 500 polyethersulfone, 10kDa, Sartorius, Germany), and samples were then reduced with 4.2 μL of 5 mM dithiothreitol (DTT) (Fluorochem, U.K.) for 40 min at room temperature and alkylated with 16.8 μL of 15 mM iodoacetamide (Sigma-Aldrich, U.K.) for 45 min in the dark at room temperature. Samples were then quenched with another 4.2 μL of 5 mM DTT, then digested with 0.4 μg of trypsin (Promega, U.K.) for 5 hours at 37°C and finally frozen. By adding 15 μL of 1 v/v% trifluoroacetic acid (TFA) (Fluorochem, U.K.), the digestion was stopped and the samples were then desalted, concentrated and purified using OMIX C18 pipette tips (Agilent Technologies, U.S.A.) with 0.1 v/v% TFA as washing solution and 50 v/v% acetonitrile (ACN) (Thermo Fisher Scientific, U.K.)/0.1 v/v% TFA as a conditioning solution. Pipette tips were prepared with two volumes of 100 μL of 0.1 v/v% TFA and washed twice with 100 μL of 50 v/v% ACN/0.1 v/v% TFA. The sample was then aspirated into the tip at least ten times to efficiently bind peptides to the absorbent membrane. Finally, two washing steps with 100μL of 0.1 v/v% TFA were performed, prior to peptides elution into 100 μL of 50 v/v% ACN/0.1 v/v% TFA. Purified peptides were left in the fume cupboard at room temperature with lids open to dry prior to their submission for LC-MS/MS analysis.

### LC/MS-MS analysis

Samples resuspended in 5 v/v% ACN/0.1 v/v% TFA were analyzed by LC-MS/MS using an Ultimate™ 3000 Rapid Separation LC (RSLC) nano LC system (Dionex Corporation, Sunnyvale, CA, USA) coupled to a Q Exactive™ Plus Hybrid Quadrupole-Orbitrap Mass Spectrometer (Thermo Fisher Scientific, Waltham, MA, U.S.A.). Peptides were separated on an EASY-Spray™ reverse phase LC Column (500 mm x 75 μm diameter (i.d.), 2 μm, Thermo Fisher Scientific, Waltham, MA, USA) using a gradient from 96 v/v% A (0.1 v/v% FA in 5 v/v% ACN) and 4 v/v% B (0.1 v/v% FA in 95 v/v% ACN) to 8 v/v%, 30 v/v% and 50% B at 14, 50, and 60 min, respectively, at a flow rate of 300 nL min-1. Acclaim™ PepMap™ 100 C18 LC Column (5 mm x 0.3 mm i.d., 5 μm, 100 Å, Thermo Fisher Scientific) was used as trap column at a flow rate of 25 μL min-1 kept at 45 °C. The LC separation was followed by a cleaning cycle with an additional 15 min of column equilibration time. Then, peptide ions were analyzed in full scan MS scanning mode at 35,000 MS resolution with an automatic gain control (AGC) of 1e6, injection time of 200 ms and scan range of 375-1,400 m/z. The top ten most abundant ions were selected for data-dependent MS/MS analysis with a normalized collision energy (NCE) level of 30 performed at 17,500 MS resolution with an AGC of 1e5 and maximum injection time of 100 ms. The isolation window was set to 2.0 m/z, with an underfilled ratio of 0.4%, dynamic exclusion was employed; thus, one repeat scan (i.e., two MS/MS scans in total) was acquired in a 45 s repeat duration with the precursor being excluded for the subsequent 45 s.

### Data analysis and statistical analysis

Peptide mass spectra were then searched against the SwissProt_2019_11 database (selected for Homo sapiens, unknown version, 20368 entries) using the Mascot search engine (version 2.5.1; www.matrixscience.com) for matches to primary protein sequences. This search included the fixed carbamidomethyl modification of cysteine as it results from addition of DTT to proteins. Deamidation (asparagine and glutamine) and oxidation (lysine, methionine and proline) were considered as variable modifications. The enzyme was set to trypsin with a maximum of two missed cleavages allowed. Mass tolerances for precursor and fragmented ions were set at 5 ppm and 0.5 Da, respectively. It was assumed that all spectra hold either 2+ or 3+ charged precursors. Scaffold (version Scaffold_4.10.0, Proteome Software Inc., Portland, OR) was used to validate MS/MS based peptide and protein identifications. Peptide identifications were accepted if they could be established at greater than 95.0% probability to maximise the reliability of the identifications. Peptide Probabilities from Mascot were assigned by the Scaffold Local FDR algorithm and by the Peptide Prophet algorithm^58^ with Scaffold delta-mass correction. Protein identifications were accepted if they could be established at greater than 90.0% probability and contained at least 2 identified peptides, in order to filter for the most accurate matches. This resulted in having a calculated decoy FRD of 0.06% for peptides and 1.9% for proteins. Protein probabilities were assigned by the Protein Prophet algorithm^59^. Proteins that contained similar peptides and could not be differentiated based on MS/MS analysis alone were grouped to satisfy the principles of parsimony. Proteins sharing significant peptide evidence were grouped into clusters. Progenesis Qi for Proteomics (version 4.1; Nonlinear Dynamics, Newcastle, U.K.) was used to perform relative quantitation calculations using the recorded ion intensities (area under the curve, AUC) and averaging the N most abundant peptides for each protein (Hi-N method, where N=3) and protein and post-translational modifications identifications. In order to increase the reliability of the matches, peptide ions with a score of <28, which indicates identity or extensive homology (p < 0.05), were excluded from the analysis based on the Mascot evaluation of the peptide score distribution for the searched .mgf file originating from Progenesis (combining all the samples in a single experiment). To further improve the reliability of the findings we implemented an additional level of filtering, excluding proteins with a peptide count of <2. Samples were grouped together using the between-subject design scheme in Progenesis, in order to compare selected group of samples (e.g. skeletonised versus fresh bones) and to calculate ANOVA p-values and maximum fold changes accordingly. The use of three biological replicates per targeted bone sample provided a sufficiently large dataset for comparative analysis using non-parametric statistical methods such as ANOVA, posthoc pairwise comparison, and Kruskal-Wallis tests. To identify proteins of interest, proteins were flagged up in order to highlight the ones that had an ANOVA p-value ≤ 0.05 and a maximum fold change ≥ 2. Common contaminants such as keratins were excluded from the interpretation of the results. Plots were done using R studio version 1.2.5033 with packages dplyr, ggplot2, ggpubr and patchwork packages. When plotting boxplots, for data following a normal distribution student’s t test and one-way ANOVA and post-hoc pairwise comparisons were used to test mean differences, otherwise Wilcoxon rank sum test and Kruskal Wallis test with post-hoc pairwise comparisons were used. STRING software version 11.0 was used to visualize functional links between the extracted proteins ^60^. The confidence score required for showing interactions was set to “high = 0.700”. MCL clustering method was used to identify the clusters, with inflation parameter = 1.5.

## Data availability

The mass spectrometry proteomics data have been deposited to the ProteomeXchange Consortium via the PRIDE^61^ partner repository with the dataset identifier PXD019693 and 10.6019/PXD019693”.

## Reviewer account details

**Password: QkJJd6VZ**

## Acknowledgements

The authors would like to acknowledge the Royal Society for funding a Research Grant (N.P.) under grant RGS/R1/191371, the UKRI for funding part of the research through a Fellowship (N.P.) under grant MR/S032878/1, as well as the European Research Council for funding part of the research under grant 319209 and the Leiden University Fund for funding under Byvanck grant 5604/30-4-2015/Byvanck. Dr. William Cheung at the NUOmics Facility is acknowledged for conducting the LC-MS/MS runs. The authors also gratefully acknowledge the donors and their next of kin for allowing the use of donated bodies to perform this research.

## Author Contributions

H. L.M. and N.P. conceived the study and wrote the paper. Text editing by all co-authors. N.P. was the lead on all proteomics experiments. H.L.M. was the lead on the human decomposition experiments and bone sampling at FACTS. D.W. was the lead on environmental data collection at FARF. H.L.M., N.P. and D.C. contributed to the interpretation of the results. N.P. and S.S. executed protein extraction, N.P. and E.S. conducted data analysis and F.S. and H.M. contributed to data analysis and to the creation of graphical outputs tables and supplementary data.

## Supplementary information

Supplementary Table 1:

Weather data over the course of the experiment period. Data were recorded by two HOBO Micro Station data loggers located on FARF at 30-minute intervals.

Supplementary Table 2:

ADD data during collection of the bone samples and samples taken for proteomics.

Supplementary Data 1:

List of 133 Quantifiable proteins extracted from Progenesis and their localisation and binding site according to Uniprot.

Supplementary Data 2:

Top 20 enriched GO terms for Biological Process (first sheet), Molecular Function (second sheet) and Cellular Component (third sheet) of the proteins extracted from all the samples.

Supplementary Data 3:

List of proteins identified in the four bone types (iliac fresh and skeletonized, tibia fresh and skeletonized).

Supplementary Data 4:

Relative abundances of proteins extracted from all samples only and grouped by bone type (iliac fresh, tibia fresh, iliac skeletonized, tibia skeletonized).

Supplementary Data 5:

List of proteins more abundant in iliac fresh samples than in iliac skeletonized samples (first sheet) and more abundant in tibia fresh samples than in tibia skeletonized samples (second sheet).

Supplementary Data 6:

Relative abundances of proteins extracted from fresh samples only and grouped by donor (D1, D2, D3, D4).

Supplementary Data 7:

Relative abundances of proteins extracted from skeletonized samples only and grouped by deposition type (open pit vs. shallow burial).

