## Supplementary Information for "The Effects of Inter-Individual Biological Differences and Taphonomic Alteration on Human Bone Protein Profiles: Implications for the Development of PMI/AAD Estimation Methods"

**Supplementary Materials and Methods**

**Informed consent at Forensic Anthropology Center at Texas State University (FACTS)**

Whole body donations are received by the Forensic Anthropology Center at Texas State University (FACTS) for scientific research under the Texas revised Uniform Anatomical Gift Act (National Conference of Commissioners on Uniform State Laws 2009). Body donations are exclusively acquired by Texas State University through the expressed and documented willing of the donors and/or their legal next of kin. Body donations are made directly to FACTS, and donors and/or their legal next of kin are aware that donations are used for taphonomic studies. Demographic, health, and other information are provided through a questionnaire completed by the donor or legal next of kin. The body donation program complies with all legal and ethical standards associated with the use of human remains for scientific research.

**Environmental characteristics at Forensic Anthropology Research Facility (FARF)**

FARF’s terrain consists of shallow clay-rich soil overlying deposits of limestone with grassland punctuated by woodlands of primarily oak and cedar trees^1^. The region has a semi-humid climate with hot summers and moderate winters. The mean average temperature is 19.4°C (68°F), but temperatures of 32°C (90°F) or greater occur in every month but December and January. Freezing is uncommon, occurring on average 36 days per year, but low temperatures reaching the freezing mark have been reported in all but five months (May through September)^2^. Annual precipitation is 81 cm, but the region experiences frequent droughts and flash flooding. The soils in the area are primarily shallow, stony and overlay Cretaceous limestone. The body donations used in this study were placed in an area of FARF with Rumple-Comfort Association, Ungulating (RUD) type soil, which is relatively shallow and rocky, with clay top soils. This soil is relatively alkaline with high carbonate content and relatively low organic matter. Because of the high clay and rock content, the soil at FARF has relatively low permeability to air and water movement^3–5^.

**Human body donations**

Donation 1

Donation 1 is the willed body donation of a 91 year old female. Donation 1 was placed on 29 April 2015 to decompose unclothed in an open pit throughout the duration of the experiment. The skeletonized remains were retrieved on 3 December 2015. A metal cage was placed over the body to protect it from large scavengers.

Bone samples were collected on 28 April 2015 (B1A) and 3 December 2015 (B1C). The mid-shaft tibia was observed to be hard, dense and of ivory white colour during drilling of the B1A sample. The iliac crest was likewise observed to be hard, dense and of ivory white colour during drilling of the B1A sample. Both mid-shaft tibia and iliac crest B1C samples displayed discoloration of the bone surface consistent with contact with soil, and were both observed to be hard and dense during drilling.

Donation 2

Donation 2 is the willed body donation of a 67 year old female with an extensive documented history of breast cancer and metastases from carcinoid tumours of unknown location. Mastectomy of both breasts was performed at 16 and 13 years prior to death. Bowel resection was performed due to a carcinoid tumour 15 years prior to death. Metastases from carcinoid tumours occurred thereafter until death. Chemotherapy treatment was given between 2 years and 7 months prior to death. Further medical information indicates prolonged consumption of calcium lactate supplements (daily for 15 years prior to death), and possible regular consumption of probiotics throughout adulthood. Bilateral nephrostomy tubes and ureter stents were placed due to blockage associated with carcinoid tumour growth and metastases. During excavation, an ellipsoidal calcified mass measuring approximately 5 cm max. length was recovered from the lower abdominal area (likely a bladder stone). Dietary information indicates a healthy diet overall, and general avoidance of processed foods and milk during adulthood.

Donation 2 was buried unclothed in a small oval pit. Bone samples were collected and the body was buried on 7 May 2015 (B2A). The skeletonized remains were excavated on 16 and 17 August 2017 and bone samples were again collected (B2C).

The mid-shaft tibia was observed to be *very* hard, dense and of ivory white colour during drilling of the B2A sample. The iliac crest was observed to be hard, dense and of ivory white colour during drilling of the B2A sample. Both mid-shaft tibia and iliac crest B2C samples displayed some discoloration of the bone surface consistent with contact with soil, and were both observed to be very hard and dense during drilling.

Donation 3

Donation 3 is the willed body donation of a 61 year old female who underwent a hysterectomy approximately 33 years prior to death. Donation 3 was buried unclothed in a small oval pit. Bone samples were collected and the body was placed on 24 June 2015 (B3A). The skeletonized remains were excavated on 21 and 22 August 2017 and bone samples were again collected (B3C).

The mid-shaft tibia and iliac crest were observed to be of lower density than the other donations, and of cream colour during drilling of the B3A samples. Both mid-shaft tibia and iliac crest B3C samples displayed discoloration of the bone surface consistent with contact with soil, and were both observed to be of lower density than the other donations during drilling.

Donation 4

Donation 4 is the willed body donation of a 77 year old female. This body donation was placed to decompose unclothed in an open pit throughout the duration of the experiment. A metal cage was placed over the body to protect it from large scavengers. Bone samples were collected on 19 October 2015 (B4A) and the body was placed on 20 October 2015. The skeletonized remains were retrieved on 9 March 2018 and bone samples were again collected (B4C).

The mid-shaft tibia was observed to be hard, dense and of ivory white colour during drilling of the B4A sample. The iliac crest was likewise observed to be hard, dense and of ivory white colour during drilling of the B4A sample. Both mid-shaft tibia and iliac crest B4C samples displayed discoloration of the bone surface consistent with contact with soil, and were both observed to be hard and dense during drilling.

**Description of the depositions**

The burials in this study were very shallow, meaning that temperature differences between open pit placements and burials are likely to have been minimal. All donors were placed in an area with the same soil type. The two burials were immediately covered with soil, while both open pit placements accumulated soil and debris at the bottom of the pits over time, which led to the covering of the pelvic sampling area with a small amount of soil (the tibia sampling area remained exposed).

**Weather data**

| **Year** | **Month** | **Total precipitation (cm)** | **Average low (Celsius)** | **Average high (Celsius)** |
| --- | --- | --- | --- | --- |
| 2015 | April (only 29 and 30 April) | 0 | 25.00 | 8.89 |
|  | May | 34.14 | 27.93 | 19.56 |
|  | June | 6.55 | 31.46 | 20.91 |
|  | July | 1.63 | 34.44 | 22.29 |
|  | August | 1.6 | 35.81 | 22.03 |
|  | September | 9.55 | 33.67 | 20.17 |
|  | October | 19.91 | 29.46 | 15.07 |
|  | November | 3.68 | 21.33 | 10.61 |
|  | December | 5.82 | 20.00 | 5.95 |
| 2016 | January | 2.64 | 16.81 | 2.81 |
|  | February | 2.9 | 21.63 | 13.84 |
|  | March | 9.68 | 24.09 | 11.43 |
|  | April | 16.46 | 26.07 | 14.17 |
|  | May | 38.48 | 28.03 | 18.05 |
|  | June | 8.38 | 33.33 | 22.22 |
|  | July | 14.35 | 36.16 | 24.25 |
|  | August | 21.11 | 33.28 | 23.32 |
|  | September | 8.05 | 32.35 | 21.35 |
|  | October | 0.69 | 30.09 | 16.02 |
|  | November | 6.17 | 24.37 | 11.78 |
|  | December | 7.9 | 17.76 | 7.67 |
| 2017 | January | 11.18 | 20.05 | 6.20 |
|  | February | 6.76 | 24.40 | 10.99 |
|  | March | 8.46 | 25.38 | 13.80 |
|  | April | 2.92 | 27.83 | 15.22 |
|  | May | 7.54 | 30.81 | 17.69 |
|  | June | 6.22 | 34.22 | 22.85 |
|  | July | 2.69 | 37.17 | 23.82 |
|  | August | 32.99 | 23.53 | 34.53 |
|  | September | 9.68 | 19.61 | 31.96 |
|  | October | 4.19 | 13.57 | 27.20 |
|  | November | 2.92 | 11.31 | 24.09 |
|  | December | 9.27 | 4.21 | 16.38 |
| 2018 | January | 0.64 | 1.02 | 15.48 |
|  | February | 2.72 | 8.27 | 17.72 |
|  | March (only 1-9 March) | 0.25 | 10.19 | 22.90 |

**Supplementary Table 1**. Weather data over the course of the experiment period. Data were recorded by two HOBO Micro Station data loggers located on FARF at 30-minute intervals.

| **Samples** | **Samples for proteomics** | **Date of collection** | **Total ADD**  **(accumulated degree days)** | **Stage of Decay**^9^ |
| --- | --- | --- | --- | --- |
| B1A-tibia | D1_TF_A  D1_TF_B  D1_TF_C | 28-04-2015 | 0 | Fresh |
| B1A-iliac | D1_IF_A  D1_IF_B  D1_IF_C | 28-04-2015 | 0 | Fresh |
| B1C-tibia | D1_TS_A  D1_TS_B  D1_TS_C | 03-12-2015 | 5225.50 | Skeletonized |
| B1C-iliac | D1_IS_A  D1_IS_B  D1_IS_C | 03-12-2015 | 5225.50 | Skeletonized |
| B2A-tibia | D2_TF_A  D2_TF_B  D2_TF_C | 07-05-2015 | 0 | Fresh |
| B2A-iliac | D2_IF_A  D2_IF_B  D2_IF_C | 07-05-2015 | 0 | Fresh |
| B2C-tibia | D2_TS_A  D2_TS_B  D2_TS_C | 17-08-2017 | 17828.40 | Skeletonized |
| B2C-iliac | D2_IS_A  D2_IS_B  D2_IS_C | 17-08-2017 | 17828.40 | Skeletonized |
| B3A-tibia | D3_TF_A  D3_TF_B  D3_TF_C | 24-06-2015 | 0 | Fresh |
| B3A-iliac | D3_IF_A  D3_IF_B  D3_IF_C | 24-06-2015 | 0 | Fresh |
| B3C-tibia | D3_TS_A  D3_TS_B  D3_TS_C | 22-08-2017 | 16817.06 | Skeletonized |
| B3C-iliac | D3_IS_A  D3_IS_B  D3_IS_C | 22-08-2017 | 16817.06 | Skeletonized |
| B4A-tibia | D4_TF_A  D4_TF_B  D4_TF_C | 19-10-2015 | 0 | Fresh |
| B4A-iliac | D4_IF_A  D4_IF_B  D4_IF_C | 19-10-2015 | 0 | Fresh |
| B4C-tibia | D4_TS_A  D4_TS_B  D4_TS_C | 09-03-2018 | 16848.23 | Skeletonized |
| B4C-iliac | D4_IS_A  D4_IS_B  D4_IS_C | 09-03-2018 | 16848.23 | Skeletonized |

**Supplementary Table 2**. ADD data during collection of the bone samples and samples taken for proteomics.
